# Key Nucleation Stages and Associated Molecular Determinants and Processes in pH-Induced Formation of Amyloid Beta Oligomers as Revealed by High-Speed AFM

**DOI:** 10.1101/2020.10.17.343830

**Authors:** Lei Feng, Hiroki Watanabe, Paul J. Molino, Gordon G. Wallace, Son L. Phung, Takayuki Uchihashi, Michael J. Higgins

## Abstract

Non-fibrillar oligomers formed via nucleation from amyloid beta (AB) peptides are currently implicated in the neurotoxicity of Alzheimer’s disease. Thus, shedding light on the molecular mechanisms underlying their formation is important for identifying targets for drug therapies. This is however an enduring challenge due to the inability to detect AB nucleation processes in the lag phase from bulk kinetic assays, while time-course analyses using a series of peptide solutions involve the discontinuous observation of dynamic nucleation processes and pathways. In this study, by adjusting the pH of AB42 peptide samples while simultaneously imaging with high-speed atomic force microscopy (HS-AFM), we show the in-situ, continuous visualization of dynamic AB42 nucleation at the molecular level. The process reveals a pH-induced saturation regime, enabling a critical monomer substrate concentration to initiate the birth of nucleation. A number of key nucleation phases are identified, including pre-nucleation, saturation regime (mass surface adsorption), nucleation and post-nucleation growth, eventually leading to the formation of predominately oligomer species. HS-AFM observations further reveal the distinct, molecular processes associated with each nucleation phase that constitute the path-dependent formation of different AB species, namely an intial monomer “diffuse-like” surface layer, followed by the emergence of nuclei and then subsequent formation and growth of new complexes and oligomers. In particular, the ability of individual nuclei to undergo surface diffusion and establish new complexes via binding interactions with other species encountered in the system was found to be a significant mechanism influencing the growth of oligomers. Herein, the study contributes to current AB nucleation theories by ascribing new molecular mechanisms. More generally, the knowledge gained from single molecule techniques can greatly assist in our current understanding of various biological processes of AB peptide such as nucleation, growth, aggregation and their related kinetic pathways.

## Introduction

Currently, there are two main nucleation mechanisms involving the formation of oligomers from Aβ monomers [1]. The primary (homogeneous) nucleation involves the growth of individual nuclei through only monomer addition in bulk solution without ‘seeds’ [2], while a more recently discovered secondary nucleation occurs on the surface of an existing amyloid structure, such as fibrils, which “catalyze” the growth of new nuclei from the monomer interactions [3]. Similarly, nucleation can be initiated by an artificial substrate, e.g. air-water interface, though this remains a primary (heterogeneous) nucleation mechanism as there is only a dependency on the monomer concentration, i.e. the substrate does not have autocatalytic activity [4]. These models are normally applied to kinetic aggregation/growth assays and importantly provide a link between the macroscopic observations, such as the typical sigmoid-like amyloid growth, and the proposed underlying molecular mechanisms. However, molecular modelling and single molecule techniques are increasingly revealing a diversity of oligomeric structures and single molecule nucleation/aggregation dynamics that are not easily detected in bulk assays and ostensibly more complex than depicted in nucleation models, which are primarily lacking details on the underlying molecular events.

One such single molecule technique, High-Speed Atomic Fore Microscopy (HS-AFM), is enabling nanometer resolution imaging on millisecond timescales to observe structural-dynamics of single Aβ monomers, oligomers and fibrils. In particular, HS-AFM studies are implementing approaches such as injecting bio-reagents, adjusting salt or monomer concentration and changing temperature on the “fly” during imaging to induce real-time dynamic changes in Aβ, primarily with a focus on fibril growth [9, 19–21]. During the time-course of imaging fibrils, some studies have observed the preceding formation of oligomers on the substrate. For amylin1-37 peptide solutions, an initial surface covering of 1-2 nm particles appears after 30 sec followed by an increase in particle size over several minutes [20]. More specifically for Aβ_42_ in NaCl solutions, small particles referred to as nuclei appear over the course of several minutes in samples seeded with either low (LMW) or high molecular weight (HMW) Aβ_42_ fractions, i.e. comprising monomers/small oligomers and high-order oligomers, respectively [21]. The LMW go on to directly form fibril seeds, while the HMW first dissociate into smaller oligomers prior to initiating fibril seeds. Whilst these observations show aspects of nucleation, a seemingly greater focus on the growth phase mechanisms of fibril formation precludes monitoring of the early nucleation (lag phase) processes in more detail.

Herein, we use HS-AFM combined with changes in pH to induce rapid Aβ_42_ nucleation on a substrate, enabling continuous monitoring and identification of key stages and their associated molecular mechanisms, including pre-nucleation, saturation regime (surface adsorption), nucleation and post-nucleation growth. For each stage, the molecular mechanisms involving different Aβ_42_ species and their dynamic interactions are described from the HS-AFM observations and their relationship to current nucleation theories are also discussed.

## Results

### Ex-situ Method: Effect of pH on Aβ peptide solutions

The effect of pH was firstly confirmed *ex-situ* by preparing peptide solutions in microcentrifuge tubes and then observing with HS-AFM. At neutral pH 7.4 three main types of oligomers/aggregate were immediately apparent on the mica surface (Figure 1B and Movie 1) and identified in histogram analysis of their individual size dimensions (calculated area and height) showing 3 peak distributions for area (nm^2^) (Fig. 1B, middle) and 2 peak distributions for height (nm) (Fig. 1B, right). The most abundant oligomer had area of 170 nm^2^ and a second oligomer was larger with area of 410 nm^2^, with both correlating to a height of 3.2 nm. A larger species had an area of 645 nm^2^ with height of 7.7nm. These observations reproduce our previous findings using HS-AFM [5] (with all dimension values to within 5-10 %) and indicate the existence of oligomers in samples prepared from Aβ_42_ peptide solutions. The latter is not uncommon and likely due to either their rapid formation in solution [6], or incomplete disaggregation of the lypholized peptide [7]. Similarly to our previous study [6], a few smaller species appear transiently, i.e. only observed in ~ 2 consecutive frames before leaving the scan area, and show an indistinct morphology likely owing to artefact when tracking fast moving objects [8]. Recently, HS-AFM imaging at 0.05 sec intervals of fast diffusing yeast prion Sup35 NM monomers show similar difficulty in delineating an accurate morphology [9]. Eventually the prion Sup35 NM monomers become ‘invisible’ at slower imaging intervals of > 0.2 secs. These findings may explain why conventional AFM often captures oligomeric structures though seemingly fails to observe monomers despite their high concentration in the sample solution, presumably due to their diffusion on timescales faster than the imaging and/or associated lack of interaction with the mica surface.

**Figure 1.**
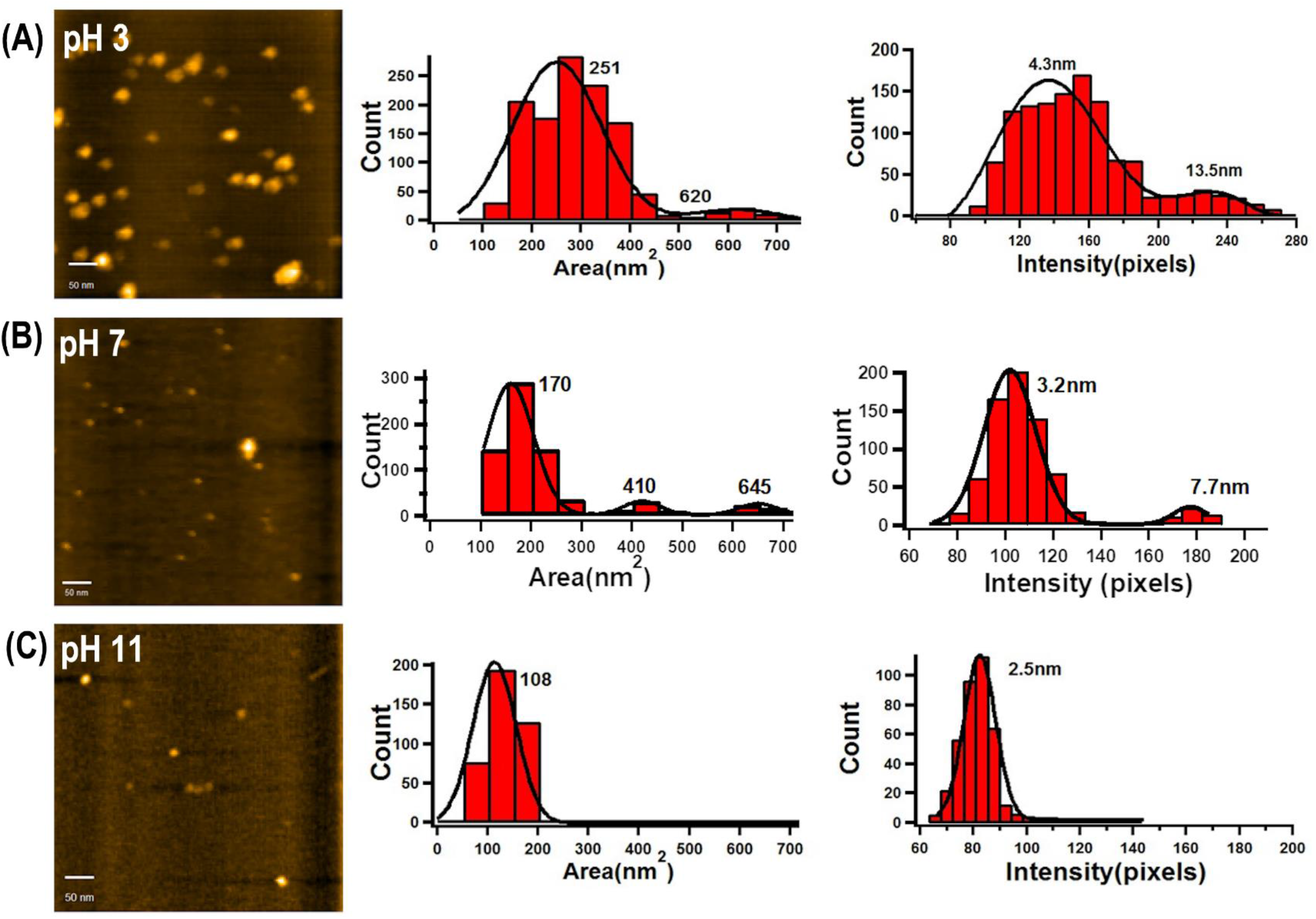
Morphology and size dimensions of Aβ_42_ oligomers prepared ex-situ in different pH solutions. (A-C) HS-AFM image of Aβ_42_ oligomers deposited on the mica surface from different pH 3 (A), 7.4 (B) and 11 (C) solutions. The 20 μg/ml peptide solutions were incubated for 1 hour at room temperature. Images obtained from HS-AFM movie with scan size 500 nm x 500nm and image rate @ 1 frame/sec. (Middle and Right) Corresponding histogram analysis of size dimensions from the calculated area in nm^2^ (Middle) and height (Right) of individual Aβ_42_ oligomers. MATLAB software automated detection of individual oligomers in each image frame and calculation of their area and height was used for the histogram analysis.

In contrast, adjusting the 20 ug/ml (~ 10 μM) peptide solution to pH 3 with an incubation time of < 30 min produces a more homogeneous, larger oligomer with particulate morphology (Fig. 1A and Movie 2). Corresponding histograms confirms this as a broadening and shift of the main distribution to higher values of ~ 250 nm^2^ (Fig. 1A, middle) accompanied by an increase in height to 4.3 nm (Fig. 1A, right). A larger species with area of 620 nm^2^ but increased height of 13.5nm remains at lower abundance (Fig. 1A, right). The pH 3 increases the number of adsorbed species on the mica surface (~55-66 molecules/μm^2^) (Figure 1A) compared to pH 7 (~30-40 molecules/μm^2^) (Fig. 1B) that is further evident by the overall total histogram counts (pH 3, count = 1287; pH 7, count = 727). In comparison histograms for pH 11 solutions show a complete loss of the 3 distinct area distributions, producing only one form of oligomer with a significantly smaller area and height of 108 nm^2^ and 2.5 nm, respectively (Fig. 1C and Movie 3). In addition, pH 11 shows a reduced number of adsorbed species (~10 molecules/μm^2^, count = 402). These observations are in accordance with the general consensus that acidic pH accelerates the rate of Aβ aggregation [10], as well as fibril formation [11]. Congo red assays demonstrate rapid aggregation of isoform Aβ_1-40_ at pH 5 within minutes compared to pH 7.4 that forms little or no aggregates in this timeframe [12]. Similarly, a greater abundance of Aβ_42_ aggregates forms during early incubation at pH 4-5, causing an increase in surface adsorbed species when imaging with AFM [13]. At acidic pH these effects are understood by the protonization of ionizable N-terminal residues, namely Glu and His [14], leading to increased positive charge density as confirmed by zeta potential measurements [13]. The latter may facilitate β-sheet conformation, which in turn promotes aggregation. Here, acidic conditions lead to an increase in size and number of the most abundant oligomer (Fig. 1A). In addition, little or negligible diffusion of oligomers on the mica surface at pH 3 (Movie 2) compared to pH 7 (Movie 1) suggests effects from a change in oligomer size, i.e. high molecular weight species diffuse more slowly, or surface charge (of both the peptide and mica surface), causing their stronger electrostatic interactions and inhibiting diffusion. Additional experiments to investigate the effect of incubation time (1, 2, and 4 hrs) and peptide concentration (from ~ 5 – 20 μM) at pH 7 did not show any significant effects other than a small rearrangement in dimensions of the existing oligomers at pH 7 (Supplementary Figures 1 and 2).

### In-Situ Method: Effect of pH on Aβ Peptide during HS-AFM Imaging

The above findings led to the rationale that changing the peptide sample solution to acidic pH during HS-AFM imaging could enable ‘real-time’ observation of the oligomer formation. Henceforth, the remainder of the study implemented experiments whereby HS-AFM imaging was initially performed on the peptide sample at pH 7 for several minutes until the imaging was optimized. Whilst continuing the imaging at typically 1 frame/sec, 2 μl of a concentrated HCl (1.0 M) solution (or NaOH for comparison) was pipetted into the liquid cell and further mixing of the solution was done by withdrawing and injecting solution with the same pipette. After an initial instability, resulting in loss of the image, the imaging could be resumed within ~ 20 seconds by adjusting the imaging parameters, including the set point, z-piezo position and gains.

### HS-AFM Observation of Aβ Nucleation Phases

A representative movie sequence of different time-points during these experiments is shown in Figure 2A along with the corresponding Movie 4. Four distinct phases are identified in the movies, including the i) pre-injection, ii) pre-nucleation, iii) nucleation, and iv) post-nucleation/growth (Fig. 2A) and correlate to the quantitative time-course analysis of particle count (Fig. 2B) and root mean square (rms) surface roughness (Fig. 2C). We note that all aforementioned phases define a process in the lag phase that is observed in classical sigmoid-shaped curves from bulk kinetic assays and known to correspond to nucleation prior to growth of fibrils defined by the elongation/growth phase [1]. Prior to injection with HCl, the oligomers are observable as described above for pH 7 (Fig. 2A, frame = 0 sec). Due to their existence prior to changing the pH, we refer to them as Aβ__EXISTING OLIGOMERS_. During this pre-injection phase (i) there is no significant change in particle count (Fig. 2B) and r.m.s. surface roughness (Fig. 2C). After the injection with HCl (Fig. 2A, frame = 30 sec), the perturbation due to injecting the solution causes the tip to lift from the surface, resulting in loss of imaging. The setpoint oscillation amplitude of the cantilever is re-adjusted to resume imaging of the surface typically within < 20 secs. During the subsequent period from ~ 30 – 130 sec (Fig. 2A, frames 60 sec and 130 sec), several Aβ__EXISTING OLIGOMERS_ show a gradual increase in size (Figure 2A, black arrow). There is also an emergence of new, smaller species on the mica surface. (Figure 2A, frame 60 sec, green arrows). During this period, the corresponding particle count analysis shows a fluctuation in values ranging between ~ 0-20 (Fig. 2B, pre-nucleation (ii)) believed to relate to detection of the new species and variability in their count as they diffuse in and out of the scan area. Time-course analysis of r.m.s. surface roughness shows a gradual increase from 0.95 nm to 1.03 nm (Fig. 2C, pre-nucleation (ii)), evidently due to the increasing size of Aβ__EXISTING OLIGOMERS_ and appearance of new species, which are both easily detected as a change in surface roughness against the atomically smooth mica surface. This period is defined as the pre-nucleation phase (ii) because it precedes a ‘mass nucleation event’ which is described below.

**Figure 2.**
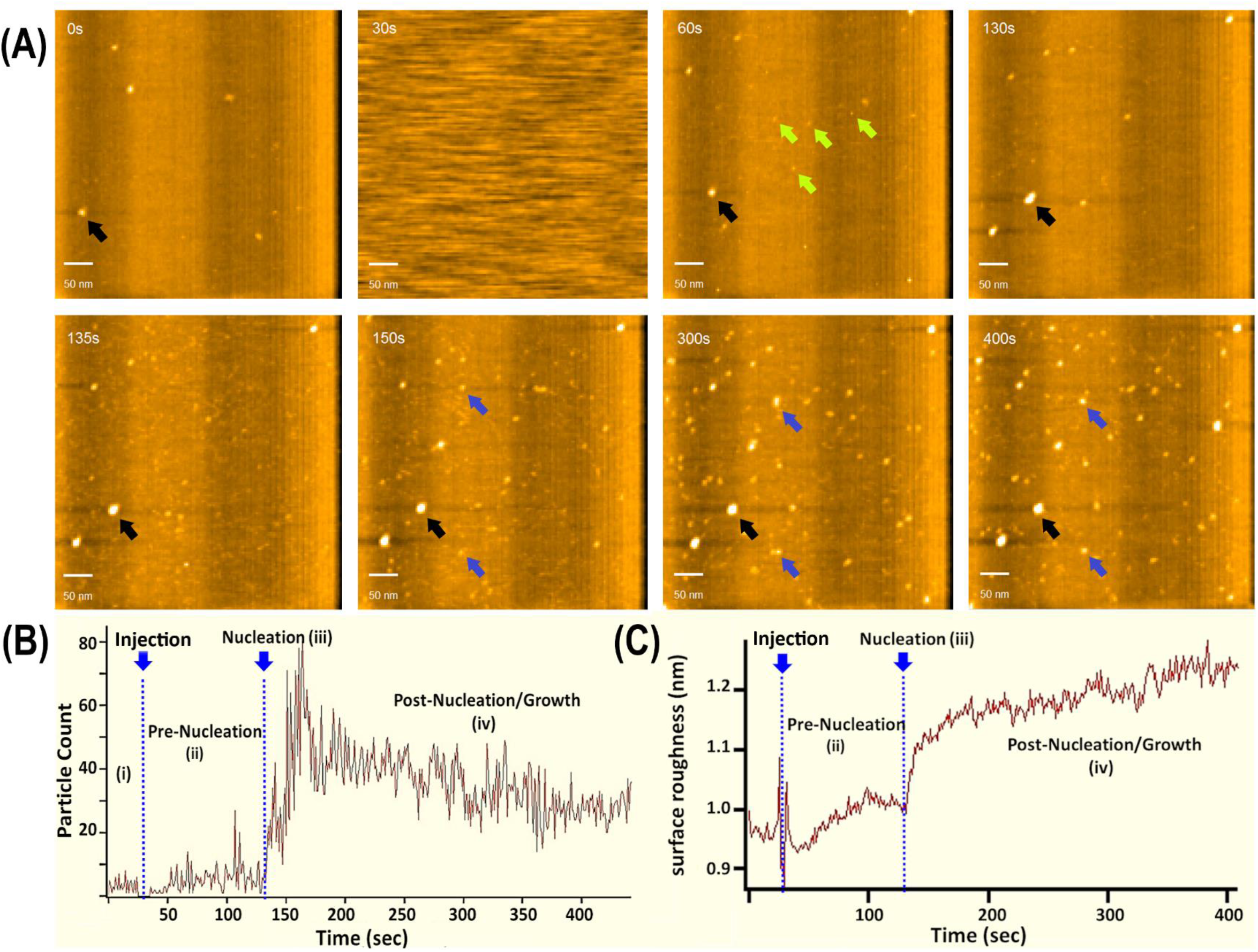
Movie sequence and time-course analysis of in-situ Aβ_42_ nucleation process after injection of HCl acid solution into 20 μg/ml peptide in PBS (pH 7.4) sample on mica surface during HS-AFM imaging. (A) HS-AFM movie sequence with scan size = 500 nm x 500 nm and image rate @ 1 frame/sec. (0 sec) Before injection of HCl solution showing Aβ oligomers on the mica surface in PBS (pH 7.4) solution. (30 sec) After injection of HCL causing loss of image. (60 and 130 sec) At pH 3 in pre-nucleation phase showing growth of existing oligomer (black arrow) from previous image frame at 0 sec and appearance of new nuclei (green arrows). (135 sec) Mass adsorption event showing rapid appearance of diffuse species covering mica surface at onset of nucleation phase. (150 sec) Nucleation phase indicating new nuclei formed during this phase (blue arrows) and the same existing oligomer (black arrow). (300 and 400 sec) Post-nucleation/growth phase showing the continued growth of the same new nuclei (blue arrows) and existing oligomer (black arrow). (B and C) Time-course analysis of (A) particle count and (B) surface roughness analysed for each image frame during all phases of the nucleation process detailed in (A). MATLAB software automated detection of individual Aβ_42_ species in each image frame was used for analysis of particle count. Igor Pro (Wavemetrics) was used for image analysis of surface roughness.

### Mass Surface Adsorption, Nucleation Phase, Post-Nucleation/Growth Phase

At a time point of ~135 sec, a significant event involving the rapid appearance of numerous smaller species is observed (Fig. 2A, frame = 135 sec). This event occurs within seconds, manifesting as a highly dynamic, diffuse species that lacks the distinct particle morphology of the Aβ__EXISTING OLIGOMERS_ and forms a covering on the mica surface (Fig. 2A, frame = 135 sec). From within this diffuse species (referred to Aβ__DIFFUSE SPECIES_), a new species of the particle-type morphology (referred to as Aβ__NUCLEI_) forms shortly afterwards, continuing to grow in size and is more clearly observable at later time-points in this phase (Fig. 2a, frame = 150 sec, blue arrows). Zoomed in images and movies (Movie 5 and 6) capture the rapid appearance of the Aβ__DIFFUSE SPECIES_ covering the mica (Fig. 3A) and nascency of the Aβ__NUCLEI_, which are initially resolved as low height, indistinct species that can diffuse on the surface and subsequently increase in size (Fig 3B, light blue arrows). As they reach larger a size, the Aβ__NUCLEI_ eventually show a cessation of surface diffusion. Timecourse analysis of particle count (Fig. 2B, nucleation (iii)) and rms surface roughness (Fig. 2C, nucleation (iii)) at this time-point (135 sec) show a concomitant linear increase in values until ceasing at ~170 sec. Hence, this period between 135 - 170 sec is referred to as the nucleation phase (iii) due to the visibly rapid onset of newly forming species, including the Aβ__DIFFUSE SPECIES_ and Aβ__NUCLEI_. Beyond this nucleation event, the Aβ__DIFFUSE SPECIES_ persists on the surface while the Aβ__NUCLEI_ can continue their growth (Fig. 2A, frames 300 sec & 400 sec, blue and black arrows, respectively). The entire period from 170 - 400 sec, referred to as post-nucleation/growth phase (iv), shows a slower rate of increasing rms surface roughness (Fig. 2C, post-nucleation/growth (iv)) though an unexpected decrease in particle count (Fig. 2B, post-nucleation/growth (iv)). This decrease is attributed to a decrease in the number of newly forming Aβ__NUCLEI_ but also the coalescence of species, which is described below.

**Figure 3.**
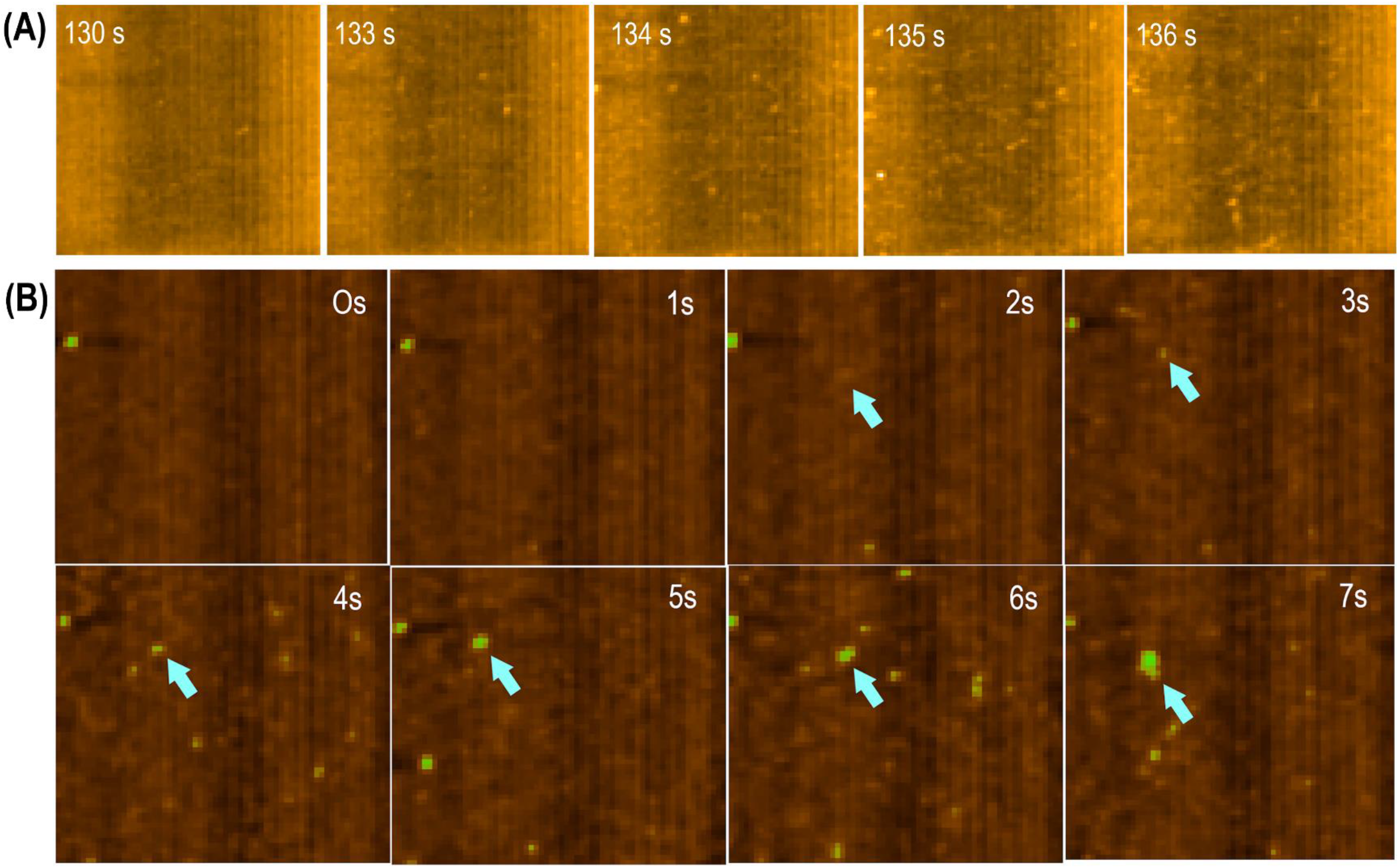
(A) Movie sequence showing mass adsorption event of diffuse species on mica surface at onset of nucleation phase. The event occurs rapidly within seconds and results in covering of the mica surface by species with indistinct morphology. Scan size 250 nm x 250 nm and image rate of 1 frame/sec (B) Movie sequence showing subsequent nucleation event indicated by the “birth” of a single nuclei. First observation is at 2 sec of low height, indistinct species (blue arrow) that appears to show diffusion to position of nuclei in next image frame. From 3 – 7 secs, the nuclei (blue arrows) shows some diffusion and continues to grow in size. Scan size 220 nm x 200 nm and @ 1frame/sec.

Time-course analysis of particle height and average area reveal further details on the different phases (Supplementary Figure 3). During the pre-injection (i) and pre-nucleation (ii) phases, particle heights of ~ 3 nm corresponding to the Aβ__EXISTING OLIGOMERS_ are observed. This rapidly increases to ~ 4 nm during the nucleation (iii) phase, taking into consideration that this value includes newly forming Aβ__NUCLEI_. Subsequently, in the post-nucleation/growth phase (iv), the height increase proceeds more slowly (Supplementary Figure. 3A). Time-course analysis of average area shows that during the pre-injection (i) and pre-nucleation phase (ii) a value of ~180 nm^2^ correlates to the Aβ__EXISTING OLIGOMERS_. The area then decreases during the nucleation phase (iii) before levelling out to ~120-140 nm^2^ in post-nucleation/growth phase (iv) (Supplementary Figure. 3B). This decrease is explained by the rapid onset of newly formed Aβ__NUCLEI_ that have a smaller area and lower the average value calculated in each image. For comparison, experiments in the alkaline condition at pH 11 by injecting the sample solution with NaoH show no changes in the samples and is confirmed by time-course analysis of particle count and rms surface roughness (Supplementary Figure 4 and Movie 7).

### Single Molecule Aβ Interactions and Complex Formation

The nucleation and post-nucleation phases were marked by the occurrence of physical interactions between individual species, typically resulting in the formation of a new complex (Movie 8). This was driven by diffusion of the Aβ__NUCLEI_ and Aβ__DIFFUSE SPECIES_, enabling an encounter with another species. When in close proximity, two individual species could come into contact to form a stable complex (Fig. 4), which we have previously observed and analyzed in-depth for interactions between the Aβ__EXISTING OLIGOMERS_ at pH 7 [5]. The binding between Aβ__NUCLEI_ is commonly observed (Fig. 4B), while both Aβ__NUCLEI_ and Aβ__DIFFUSE SPECIES_ could bind to more immobile, larger Aβ__EXISTING OLIGOMERS_ (Fig. 4C). Although more difficult to observe, the binding of fast diffusing Aβ__DIFFUSE SPECIES_ is discernible (Fig. 4A). Furthermore, an individual species could undergo multiple, continuous interactions, leading to an observable increase in size upon forming a new stable complex.

**Figure 4.**
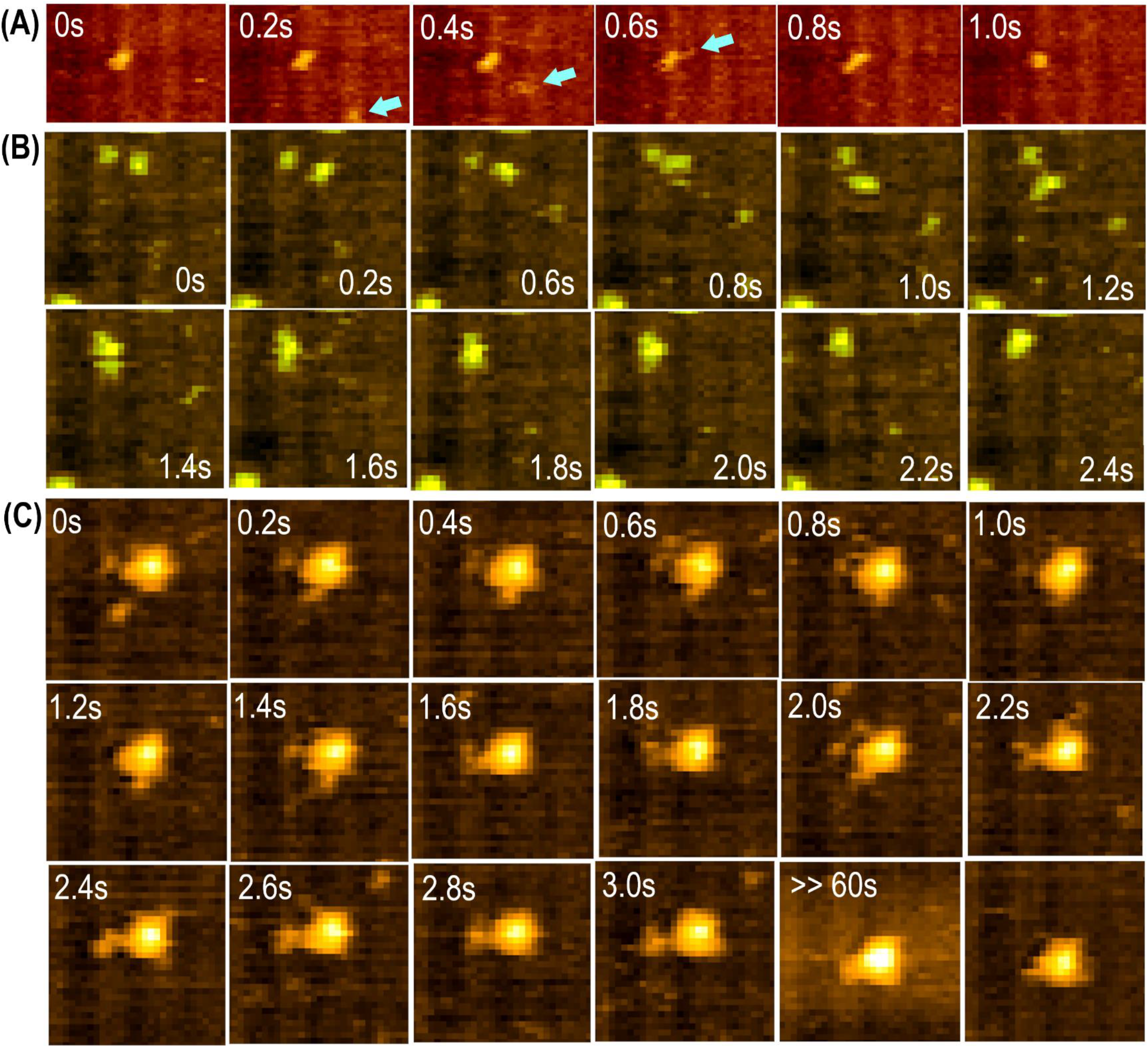
(A) Movie sequence showing the binding interaction between a single nuclei and diffuse species exhibiting fast diffusion. The diffuse species appears at 0.2 sec and then binds to the nuclei at 0.6 sec, observed as a fine appendage (blue arrow). At 0.8 and 1.0 sec, the new complex is formed, i.e. the diffuse species remains bound, and undergoes a conformational change. Scan size 90 nm x 60 nm and image rate @ 5 frames/sec (B) Movie sequence showing binding of two single nuclei in close proximity to each other. From 0 – 1.2 sec both nuclei show conformational changes and limited diffusion. At 1.4 sec the nuclei come into contact, subsequently appearing to establish their binding from 1.6 – 2.0 sec, until a stable complex is formed at 2.2 – 2.4 sec. Scan size 60 nm x 65 nm and image rate @ 5 frames/sec. (C) Movie sequence showing larger existing oligomer participating in binding of single nuclei from 0 – 1 sec (binding to bottom of oligomer) and 2.4 sec – >> 60 sec (binding to left of oligomer) and binding of multiple species at the one time (1.4 and 2.2 sec). Scan size 60 nm x 65 nm and image rate @ 5 frames/sec.

### Growth Rate of Single Molecule Aβ Species

Movie sequences of representative individual Aβ__EXISTING OLIGOMERS_ (labeled as number 1, 2, 3 in Supp Fig.5 and) and Aβ__NUCLEI_ (labeled 4, 5, 6 in Supp Fig.5) reveal varying growth rates in height and/or area (Supplementary Fig. 5). For example, species 3 and 6 show the greatest increase in size dimensions, with corresponding time-course analysis of the same six individual species given in Figure 5. For Aβ__EXISTING OLIGOMERS_ the increase in area (Fig. 5A) and height (Fig. 5B) for species 3 occurs in a linear fashion, mostly in the pre-nucleation phase and continues into nucleation phase before leveling out. From the linear slope region, species 3 gives area and height growth rates of ~1.5 nm^2^/sec and ~0.07 nm/sec, respectively. In contrast species 1 and 2 show no significant change in area and height (Fig. 5A and B). In addition, the profiles of the three species show fluctuations in values that are difficult to rationalize as changes in growth. These are instead attributable to conformational changes of species and explains the unusual change in area values, i.e. decrease for species 3, opposite rise-fall for species 2, and sudden later increase for species 1 (Fig. 5A). This is supported by movies confirming many of the Aβ__EXISTING OLIGOMERS_ and Aβ__NUCLEI_ continuously undergo conformational changes, resulting in changes of shape factor and dimensions. The profiles of Aβ__NUCLEI_ evidently commence at a time after the onset of nucleation (Fig. 5C and D). At ~200 sec, species 6 (black trace) shows a linear increase in area while species 4 (red trace) and 5 (blue trace) show no change apart from fluctuations in values as described above (Fig. 5C). For the height commencing at ~150 sec, all species show an initial increase of 0.5 – 1.0 nm (Fig. 5D). Following this, species 4 and 5 show no further change while species 6 initiates a further increase in height at ~200 sec that is concomitant with the increase in area occurring at the same time in Fig. 5C.

**Figure 5.**
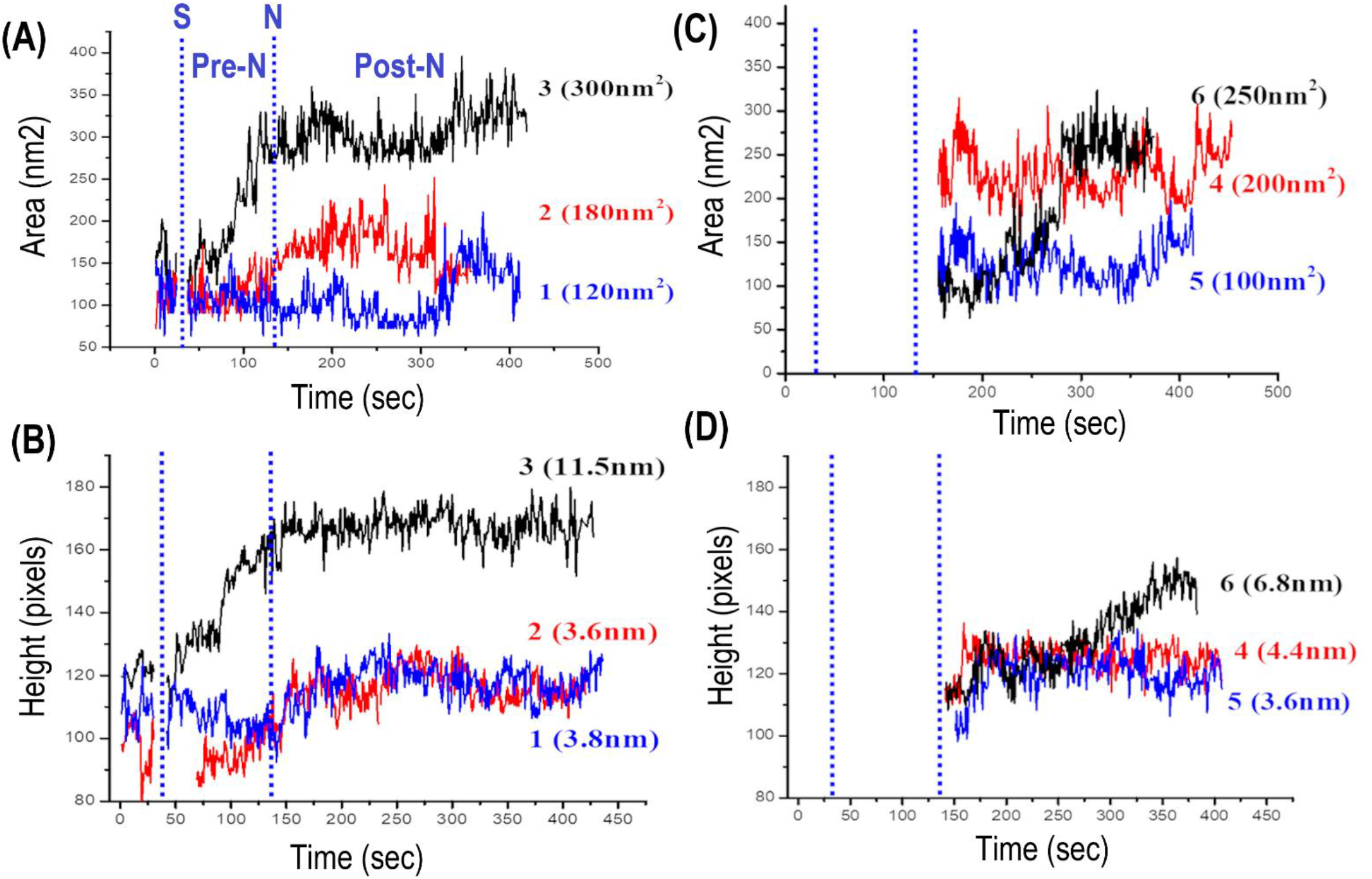
Time-course analysis of calculated area and height of representative single existing oligomers labelled 1, 2, 3 and new nuclei labelled 4, 5, 6, corresponding to images in Supplementary Figure 5. Dashed lines represent time of injection (I) and onset of nucleation (N) with pre-nucleation (Pre-N) and post-nucleation/growth (Post-N) phases. For existing oligomers, profiles of area (A) and height (B) commence prior to injection. Species 3 shows a significant increase in both area and height during pre-nucleation phase while other species 1 and 2 show no observable growth. For new nuclei, profiles of area (C) and height (D) commence after onset of nucleation (~150 sec). Species 6 shows an increase in area occurring from 200 – 300 sec. All species show an initial, small increase in height. Species 6 shows a further increase in height while species 4 and 5 show no further changes.

## Discussion

Herein, we discuss the nucleation process observed at acidic pH in the HS-AFM movies, including interpreting the key stages and their associated molecular mechanisms. Prior to injection of HCl, oligomers are present on the mica surface at pH 7. Namely, they have already formed in the sample solution prior to adsorption on the mica. These oligomers have previously been shown to be highly dynamic and capable of binding interactions to form stable complexes, particularly the larger aggregates [5, 15]. Conversely, apart from a few suspected transient species on the mica, there is no clear indication of monomers despite being present at higher concentrations in the sample, indicating they are present in the bulk solution.

After injecting with HCL in a pre-nucleation phase, the growth of existing oligomers proceeds linearly, revealing they are able to resume growth as a surface adsorbed species. At the same time, there is an emergence of new smaller species, representing the first indication of nucleation. The pH induced effect driving this oligomer re-growth and formation of new nuclei is understood by protonation of the monomer N-terminal residues [14], increasing positive charge and β-sheet content to facilitate the monomer interactions as mentioned above. As no monomers are clearly visible on the surface, the processes are assumed to be driven by the addition of positively charged monomers that diffuse from the bulk to within proximity of the mica surface. It is also possible that a supply of monomers reside at the liquid-solid interface, exhibiting lateral surface diffusion on timescales faster than the HS-AFM imaging. The existence of this pre-nucleation phase may represent the time delay in homogenous mixing of the injected sample solution and lowering of pH necessary to initiate the subsequent phases.

The main nucleation phase is marked by the onset of a mass adsorption event. This is the rapid appearance of a higher density, faster diffusing adsorbed species that largely covers the mica surface (Fig. 3A). Coinciding with this event is the formation of new nuclei (Fig. 3B). For the mass adsorption, it is interpreted that while the monomer charge is positive below its isoelectric point (i.e.p. = 5.5) the mica surface may remain weakly negative with a low i.e.p. of 3 - 3.5. Thus, a preferential electrostatic attractive interaction facilitates the mass adsorption. Critically, this surface adsorbed species, presumably the monomer or ‘clusters or assemblies’ thereof, is the source of the nuclei and their mass adsorption enables a critical concentration to initiate their nucleation. It is apparent that the latter is achieved rapidly within seconds due to the conditions favouring adsorption and confinement of monomers on a surface. This observation can be described by a saturation regime where secondary nucleation (e.g. on a fibril) may saturate or show little dependence on monomer concentration in solution when there is a high monomer concentration on the substrate [16]. This has previously been shown for Aβ_42_ upon reduced electrostatic repulsion by salt screening [17] and pH [18], to drive surface adsorption and increase in monomer substrate concentration. Furthermore, these observations imply that secondary nucleation is not a single step reaction and envisaged to involve a number of rate-limiting steps, e.g monomer-substrate association, nucleation and detachment [16].

During the nucleation phase, another key step in the process is that upon their formation, the nuclei are able to diffuse on the mica surface. This allows for subsequent encounters of the different individual species, leading to a multitude of binding interactions and formation of new complexes. For example, a newly formed nuclei can form new complexes with other nuclei or existing oligomers (Fig. 4B and C) and then those new complexes can continue to interact with other individual species. These highly pervasive interactions occur throughout the nucleation and post-nucleation phases, leading to growth akin to ‘coalescence’ of individual oligomers. in addition, the faster diffusing, diffuse species, presumably the monomers, also appear to continuously interact and bind, forming complexes with nuclei or oligomers (Fig. 4A and C) though these interactions are less easily resolved.

## Conclusion

In conclusion, the continuous visualization of Aβ_42_ nucleation processes using HS-AFM provides explicit details on the molecular mechanisms, revealing a multitude of interactions and dynamics that as yet are only gleaned from macroscopic observations. The molecular determinants include different species types (solution monomer, surface-adsorbed monomer, nuclei (new oligomers), existing oligomers and oligomer complexes) that exhibit various dynamics (including nucleation, growth, diffusion, intramolecular interactions/binding and formation of complexes) and each play a role in a series of key stages and associated molecular processes. This is summarized in Figure 6, which sheds new light on current nucleation models by ascribing the molecular details. While the nucleation on the mica surface is by definition, primary (heterogeneous) nucleation, the molecular mechanisms uncovered are applicable to secondary nucleation that is also facilitated by a substrate. The use of HS-AFM is coming of age and will continue to resolve the molecular domain of Aβ_42_ peptides and their oligomer forms, including the exciting prospect of elucidating their dynamic interactions with other biological molecules or drugs.

**Figure 6.**
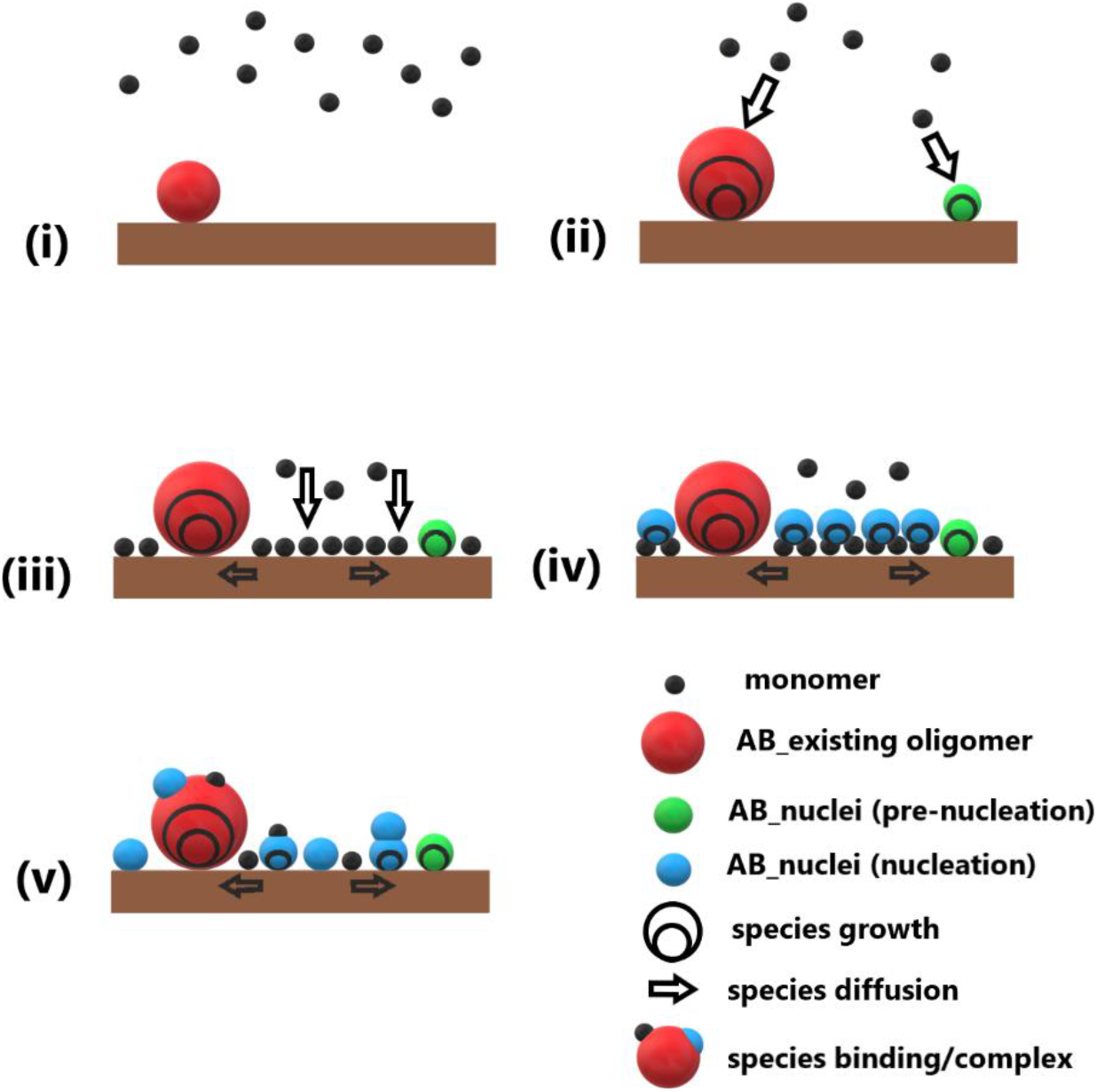
Schematic of nucleation stages and associated molecular mechanisms. (i) In the pre-injection phase at pH 7.4 (i) existing oligomers are on the mica surface and monomers are in bulk solution. (ii) After injection of the acidic solution at pH 3 the pre-nucleation involves the growth of existing oligomers and formation of nuclei through addition of positively charged (protonated) monomers diffusing in close proximity to the surface. (iii) The change in pH drives preferential surface adsorption and increase in monomer substrate concentration (‘mass adsorption event’). (iii) The critical substrate monomer concentration initiates the nucleation of nuclei (new oligomers). (iv) The surface diffusion of monomers and nuclei enables their interactions, binding and formation of new complexes, contributing to further growth of individual species.

---

**Supplementary Figure 1.**
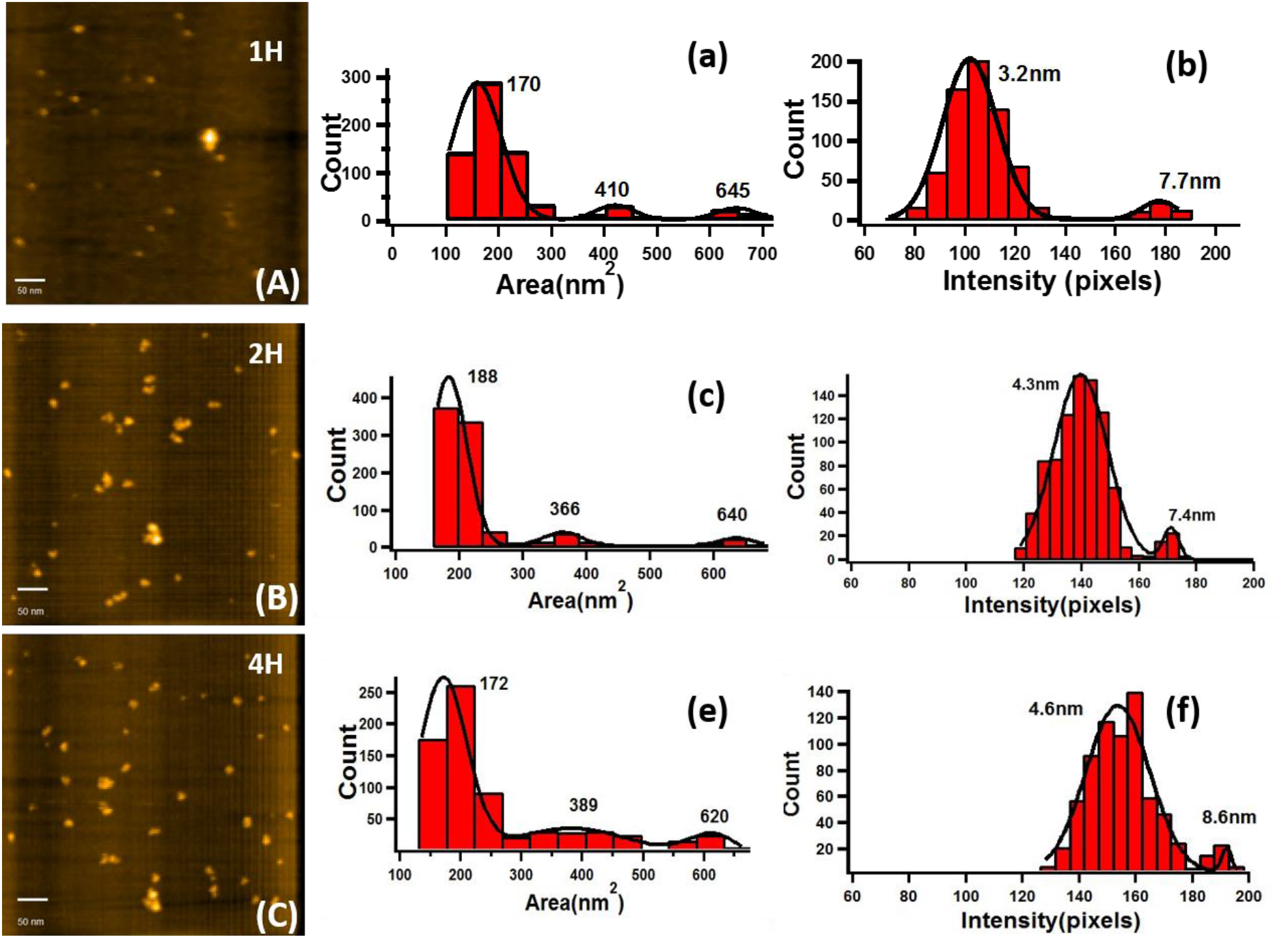
Morphology and size dimensions of Aβ_42_ oligomers prepared ex-situ using different incubation times of 1, 2 and 4 hours. (A-C) HS-AFM images of Aβ_42_ oligomers deposited on the mica surface from sample solutions incubated for 1 (A), 2 (B) and 4 (C) hours. Scan size 500 nm x 500nm and frame rate @ 1frame/sec. The 20 μg/ml peptide solutions were incubated in PBS (pH 7.4) at room temperature. (Middle and Right) Corresponding histogram analysis of size dimensions from the calculated area in nm^2^ (Middle) and height (Right) of individual Aβ_42_ oligomers.

**Supplementary Figure 2.**
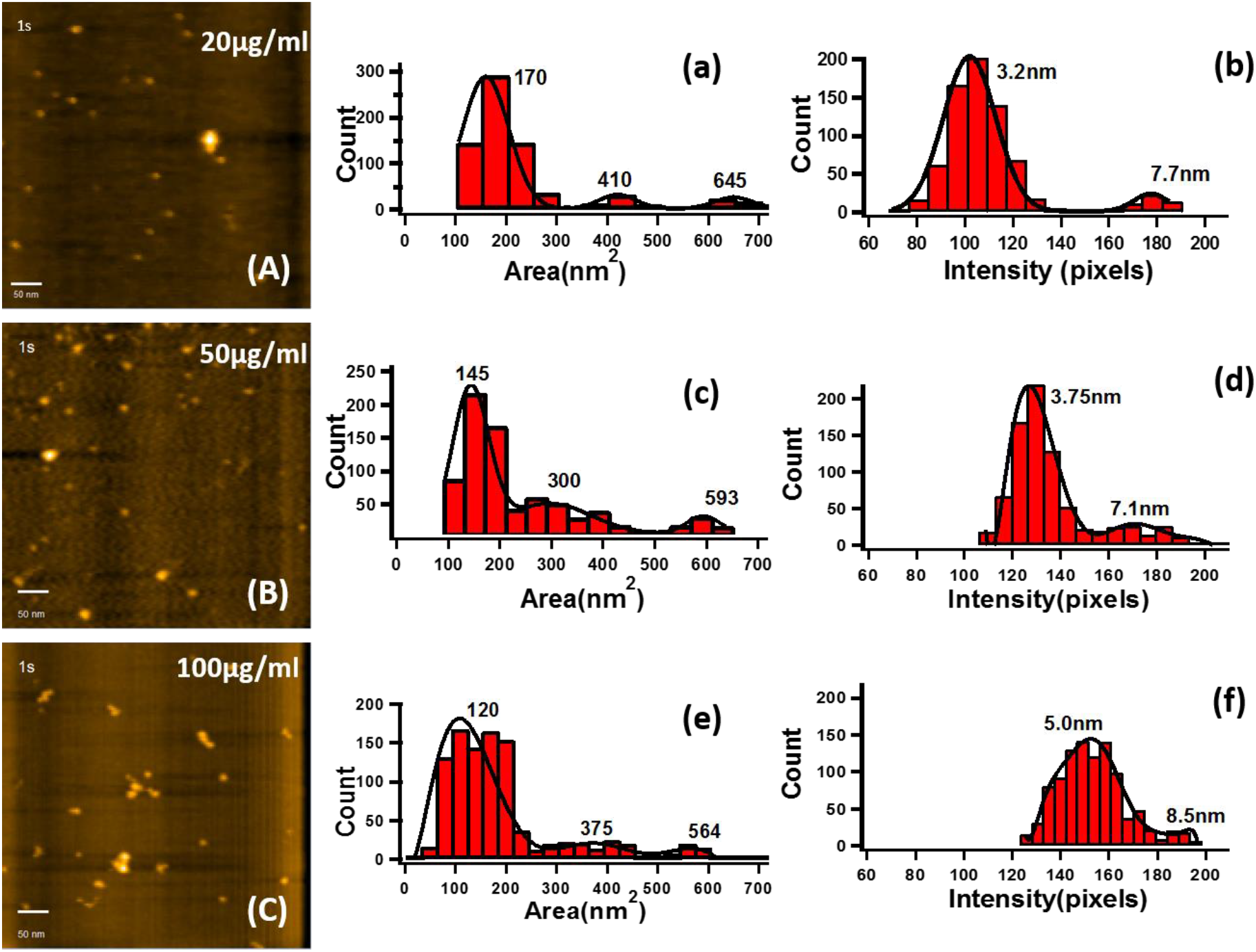
Morphology and size dimensions of Aβ_42_ oligomers prepared ex-situ using different peptide concentrations. (A-C) HS-AFM images of Aβ_42_ oligomers deposited on the mica surface from samples solution with peptide concentrations of 20 μg/ml (A), 50 μg/ml (B) and 100 μg/ml (C). Scan size 500 nm x 500nm and frame rate @ 1frame/sec. The peptide solutions were incubated in PBS (pH 7.4) for 1 hour at room temperature. (Middle and Right) Corresponding histogram analysis of size dimensions from the calculated area in nm^2^ (Middle) and height (Right) of individual Aβ_42_ oligomers.

**Supplementary Figure 3.**
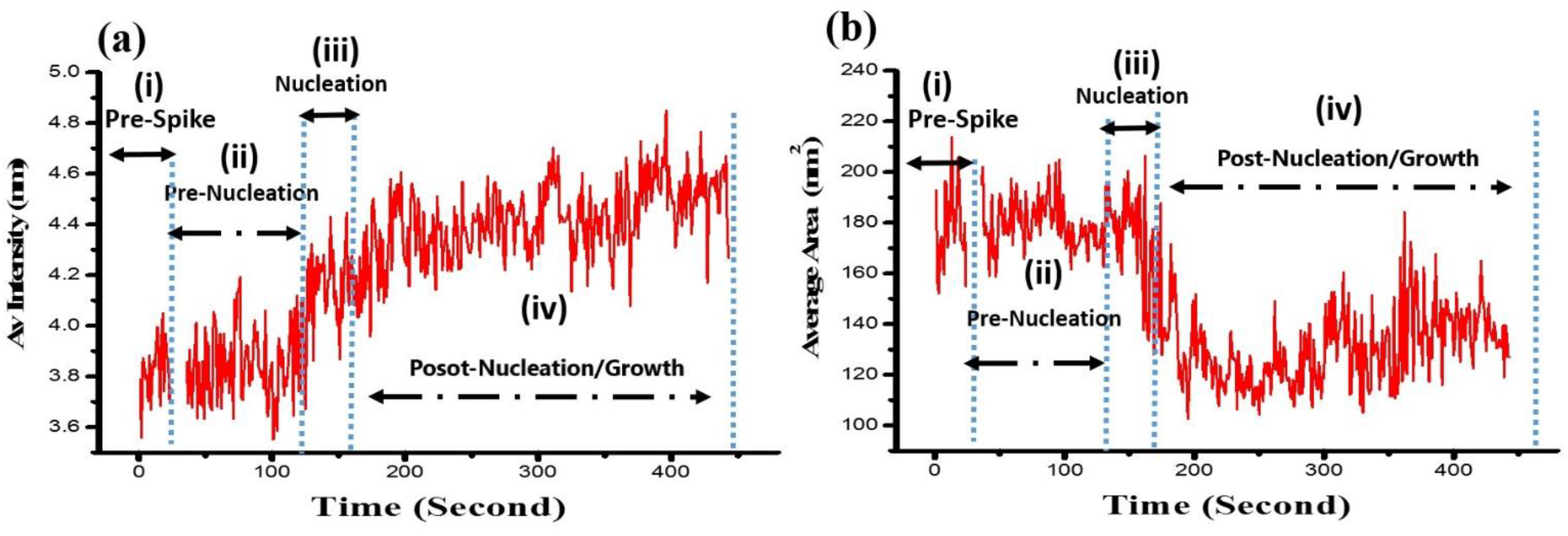
Time-course analysis of average (A) height and (B) area of Aβ_42_ species analysed for each image frame during all phases of the nucleation process detailed in Figure 3. MATLAB software automated detection of individual species in each image frame and calculation of their height and area was done for the analysis.

**Supplementary Figure 4.**
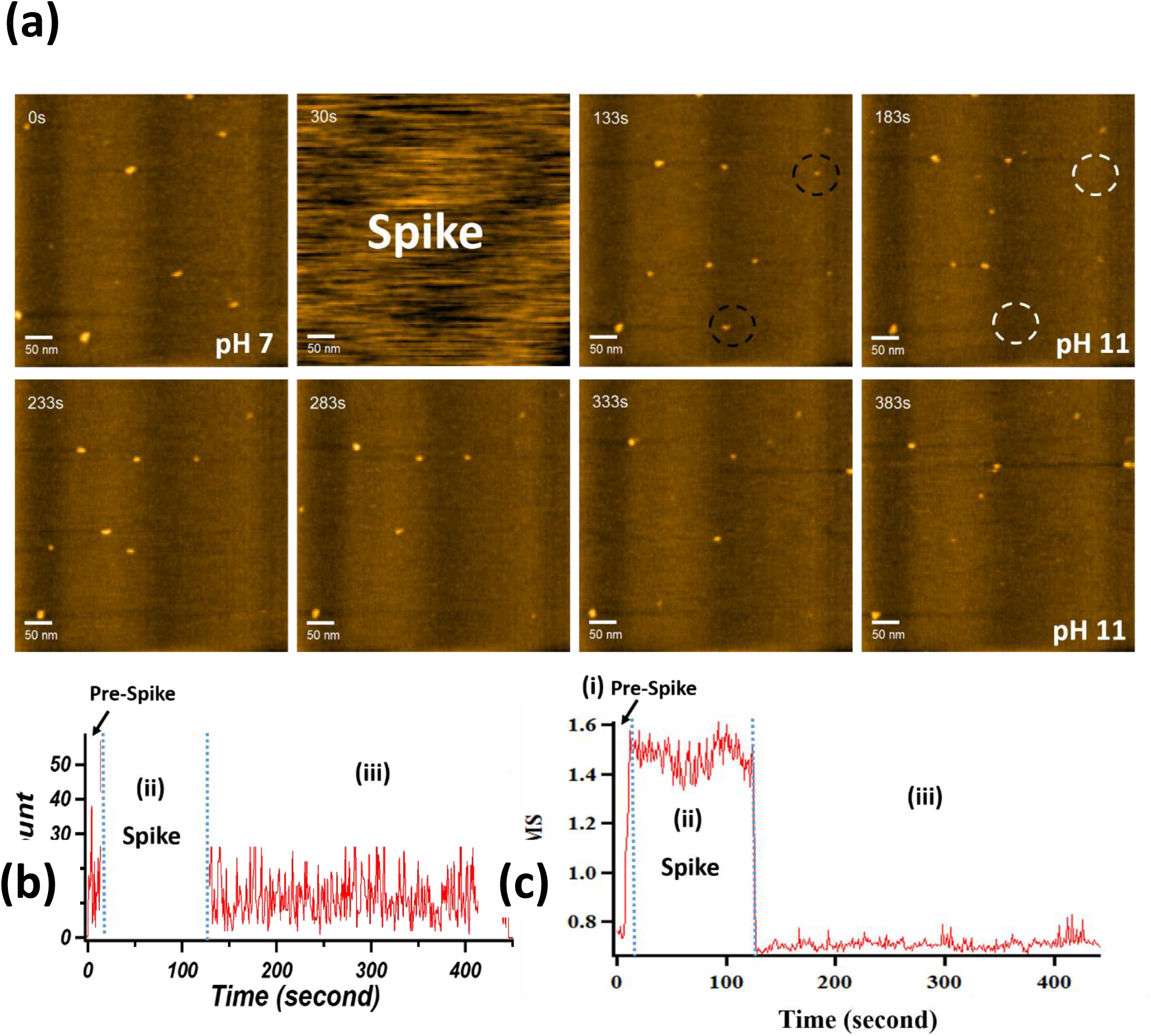
Movie sequence and time-course analysis of in-situ experiment after injection (SPIKE) of NaOH alkaline solution into 20 μg/ml peptide in PBS (pH 7.4) sample on mica surface during HS-AFM imaging. (A) HS-AFM movie sequence with scan size = 500 nm x 500 nm and image rate @ 2 frame/sec. (0 sec) Before injection of NaOH solution showing Aβ oligomers on the mica surface in PBS (pH 7.4) solution. (30 sec) After injection of NaOH causing loss of image. (133 – 383 sec) For the remainder of the sequence at pH 11, no changes or nucleation was observed. (B and C) Time-course analysis of (A) particle count and (B) surface roughness analysed for each image frame during the movie sequence in (A). No change in particle count or roughness was observed. The jump in surface roughness during the injection (SPIKE) corresponds to noise during loss of image. MATLAB software automated detection of individual Aβ_42_ species in each image frame was used for analysis of particle count. Igor Pro (Wavemetrics) was used for image analysis of surface roughness.

**Supplementary Figure 5.**
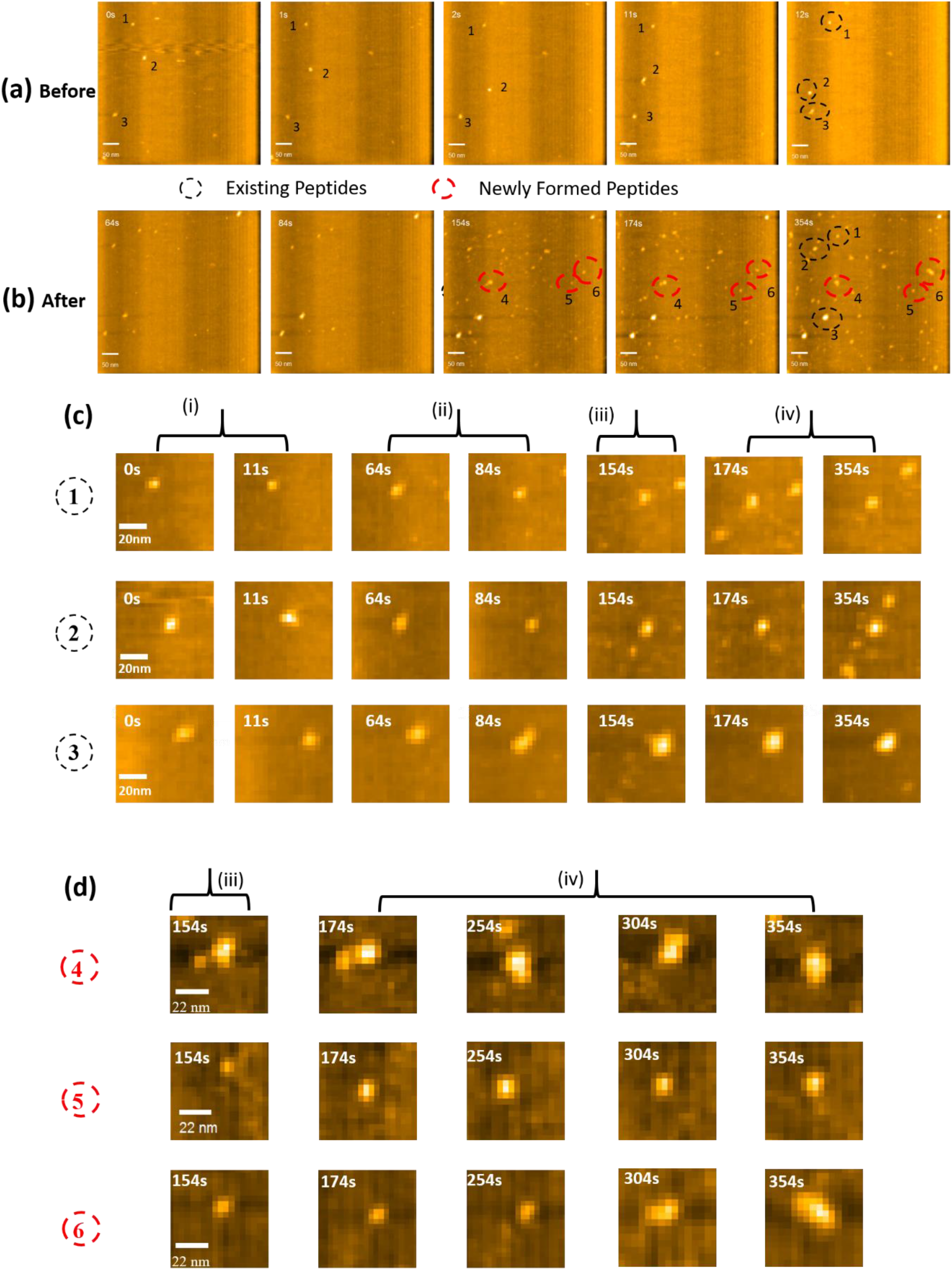
Movie sequence shown in Figure 3 with labelling of (A) individual existing oligomers (1, 2, 3) before injection and (B) new nuclei (4, 5, 6) after injection of HCl acidic solution. All species are labelled in the final image frame. Scale bar = 50 nm. Movie sequence of magnified image of each individual (B) existing oligomer 1, 2, 3 (scale bar = 20 nm) and (C) new nuclei 4, 5, 6 (scale bar = 22 nm)

## Materials and Methods

### Preparation of Aβ_42_ peptide samples

Amyloid-beta protein fragment 1-42 (Aβ_42_ peptide) was purchased as a lyophilized solid and more than 95% purity was guaranteed by HPLC from Sigma-Aldrich (High Performance Liquid Chromatography). Aβ_42_ peptide samples were prepared according to previous methods of [22, 23]. 1 mg of lyophilized solid peptides were firstly dissolved in (1,1,1,3,3,3-hexafluoro-2-propanol (HFIP) to prevent aggregation and then aliquot into 50 small microcentrifuge tubes. After 30 minutes incubation at room temperature in a chemical fume hood, the HFIP peptide solutions were allowed to evaporate for 60 minutes and the re-lyophilized peptides stored at −20°C. Lyophilized peptides were solubilized using phosphate-buffed saline (PBS) (pH 7.4) at a concentration of 20μg/ml and then vortex mixing applied for 20 seconds prior to use as the sample solution for HS-AFM imaging.

### Remodification of HS-AFM Cantilever Tips

High resonance frequency (0.8-1.2MHz) cantilevers with low spring constants (0.1-0.2 Nm^-1^) specially designed for HS-AFM were obtained from Olympus (BL-AC10DS). The silicon cantilevers consisted of a beak-like tip upon which a carbon-based tip was further deposited by electron-beam deposition (EBD). The carbon tip was sharpened by argon or oxygen plasma etching to produce a significantly smaller tip radius of ~3 nm compared to the original tip (25-100 nm) of the commercial cantilever, which enabled improved imaging stability and resolution [24, 25]. After imaging, the carbon tip could be completely removed by plasma etching and the cantilever remodified for reuse.

### High-Speed Atomic Force Microscopy

To prepare samples for imaging, 2 μl of the Aβ_42_ peptide solution was pipetted onto a 1.5 mm diameter freshly cleaved mica disc (Cat No. 7101) and incubated for 2 minutes to allow the peptides to adsorb onto the mica surface. The sample stage with mica surface was then immersed in liquid to form a final sample solution volume of around 70 μl with PBS. 2μl of fresh PBS was then added and pipetted in/out of the sample solution, and this was repeated several times to exchange the sample solution and remove excess peptides that had not adsorbed onto the mica surface. The sample was then placed into the liquid cell of the HS-AFM. HS-AFM imaging was performed using an ANDO-model (Research Institute of Biomolecule Metrology Co., Ltd., Japan) in tapping mode with high frequency, small cantilevers (BL-AC10, Olympus) remodified with the carbon-tip, as described above.

HS-AFM imaging was performed on samples prepared with different peptide concentrations (20μg/ml, 50 μg/ml or 100 μg/ml), incubation times (1h, 2h and 4h) and pH of 3, 7, and 11. Two different experimental methods were applied, including the ‘ex-situ’ and ‘in-situ’ methods. For the ex-situ method the peptide samples as a function of concentration, incubation time and pH were prepared in a fresh centrifuge tube prior to depositing the sample solution on the cleaved mica substrate for imaging. As mentioned above, after deposition of the peptide solution for 2 min, the sample solution was then exchanged with fresh PBS by pipetting and HS-AFM imaging performed using the parameters/conditions described below. In contrast, the ‘in-situ’ method involved depositing 2 μl of a 20ug/ml peptide solution on the cleaved mica substrate. This was allowed to adsorb for 2 min, followed by the rinsing process as mentioned above, with a final volume of 70 μl PBS (pH 7) used for the HS-AFM imaging. Whilst imaging a pipette was used to inject 2.0 μl of 1M concentrated HCL or NaOH into the sample solution and further mixing of the sample solution was performed by withdrawing/injecting the solution via pipetting. The imaging parameters were subsequently adjusted, including the optimization of set point, gains and z-height, to stabilize the imaging. In order to obtain the desired, a dilution factor of the HCL and NAOH solution required to achieve the pH 3 and 11 in PBS was previously determined and applied to the AFM sample solution.

During imaging, the free oscillation amplitude of the cantilever was set to ~2 nm, and the set point amplitude was kept to ~ 90% of the free amplitude. The maximum possible scanning rate is calculated as R_max_= (λ*f*)/(2WN), where λ is the amplitude, *f* is feedback bandwidth, W is the scanning size, and N is the corresponding number of scan lines. A non-electrode wide scanner with a range of 4 μm × 4 μm and a z scanner with a maximum height of 700 nm were used with typical scan speeds of 2-4 frames/sec for 500 nm scans. Higher resolution scans of 200 nm were taken at 5-10 frames/sec with 550-275 lines/sec. The imaging was performed for a typical duration of between 10 minutes and 2 hours or until it was clear that tip quality had reduced, presumably due to adhesion of peptides on the tip. For each sample condition, at least three separate imaging experiments using new tips were performed. The operation of the HS-AFM and collection of the AFM movies was obtained by modified software in Igor Pro, Wave metrics. AFM movies were either analyzed using Igor Pro or exported as an AVI. Movie file for further analysis using a MATLAB program.

### Analysis of HS-AFM Movies

The analysis of HS-AFM movies was done according to previously developed MATLAB code by our group. The MATLAB program was developed to import HS-AFM movies, separate individual frames, and segment objects (the peptides) within each frame. The program had four main stages: 1) background image estimation; 2) background subtraction and contrast enhancement; 3) image segmentation using the Otsu’s algorithm; 4) connected component labelling and object measurement. In Stage 1, the background image was estimated by applying image erosion followed by image dilation. Let *I*(*x,y*) be an input HS-AFM image, where *x* and *y* are the horizontal and vertical coordinate, respectively. To generate the eroded image *I_e_*, the pixel at location (*x,y*) was computed as the *minimum* of all pixels in a local neighbourhood of image *I*:

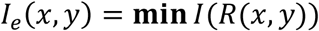

In our experiments, the local neighbourhood *R* was selected as a circular region of radius 15 centred at pixel location (*x, y*). Next, to generate the dilated image *I_b_*, the pixel at location (*x, y*) was computed as the *maximum* of all pixels in a local neighbourhood of image *I_e_*:

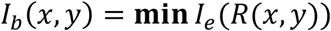

The dilated image *I_b_* (*x, y*) was considered as an estimate of the image background. In Stage 2, the background image was subtracted from the input image: *I_d_ = I – I_b_.* The difference image *I_d_* contained mostly the foreground (i.e. the molecules). To improve the contrast of the foreground and consequently the segmentation accuracy, the difference image *I_d_* was scaled linearly to the full intensity range [0, 255]. In Stage 3, the enhanced difference image was segmented using the Otsu’s algorithm. In this algorithm, a threshold is applied to separate the image into two classes: the foreground and the image background. This threshold is selected to minimize the intra-class variance and maximize the inter-class variance. In other words, this algorithm increases the similarity between pixels belonging to the same class, while decreasing the similarity between pixels belonging to different classes.

In Stage 4, connected component labelling was applied to group connected pixels into regions (molecules). For each region, several properties were measured, including width, height, bounding-box coordinates, centroid, perimeter, area, and eccentricity. In addition, the length and width of peptides were validated using the cross-section analysis in the Igor Pro software. Using the above properties, molecules were also tracked across multiple frames of the HS-AFM video sequence. We also applied image filtering to the video sequences, especially for the height measurement, to better observe the individual molecules and their dynamic interactions. Consequently, some darker regions may appear in the snapshot images and slightly affect the real height value of peptides that were adsorbed in these darker areas. In these cases, we performed a manual analysis using the line height profile in Igor (HS-AFM software) with or without the filter to calculate a correction factor for the height data.

### Statistics analysis

Values obtained from the MATLAB analysis of HS-AFM movies included the peptide area, intensity (height), width and length as described in our previous work. All data analysis and histograms were produced using Igor Pro, Wave metrics. Multi-peak fitting with Lorentzian or Gaussian functions were applied to the histograms.

